# Galectin-3 is Necessary for Selective Cathartocytosis, which Expedites the Development of Proliferative Gastric SPEM

**DOI:** 10.64898/2026.06.07.729580

**Authors:** Xiaobo Lin, Xuemei Liu, Gabriel Nicolazzi, Annie Pan, Margaret Hua, Jeffrey W. Brown

## Abstract

The expression and secretion of sulfated colonic-type mucins is a feature of high-risk metaplasias of the gastrointestinal foregut (Barrett’s esophagus, type III intestinal metaplasia of the stomach, and pancreatic intraepithelial neoplasia). Galectin-3 is a lectin that preferentially associates with galactose modified by a 3’-O-sulfate relative to its unmodified counterparts and is upregulated as the tissue transitions to high-risk metaplasia, dysplasia, and cancer. Since both galectin-3 and sulfated glycotopes are aberrantly and concurrently overexpressed in high-risk premalignant and malignant tissue transformations, we sought to investigate the role of galectin-3 in the metaplastic reaction. We found that injury induces the expression of *Lgals3* at the RNA and protein levels. Unlike cancer cell lines, we show that *in vivo* galectin-3 colocalized with sulfomucins in zymogenic granules of the gastric chief cell. Utilizing a synchronous, chemically-induced murine model that produces spasmolytic polypeptide expressing metaplasia, we found that galectin-3 facilitates cathartocytosis of the vesicles it resides in, but not organelles lacking LGALS3. Inhibition of cellular downscaling resulted in delayed expression of the metaplastic transcription factor Sox9 as well as proliferation. Here, we present a new role for galectin-3 in promoting the transition from normal, homeostatic tissue to metaplasia and our data suggest that cathartocytosis represents an unconventional secretory pathway for galectin-3, which has been a matter of controversy as galectins are not secreted via canonical pathways.

## Introduction

The ectopic expression and secretion of sulfated (colonic-type) mucins have long been associated with high-risk metaplasia and cancer throughout the gastrointestinal foregut: esophagus, stomach, and pancreas(1-6). In fact, the presence and type of acidic glycans histologically distinguishes the three types of gastric intestinal metaplasia and correlates with the propensity for oncogenic transformation (7). Specifically, the epithelial cells in complete (type I) metaplasia are devoid of acidic mucins and are considered low risk lesions. In contrast, incomplete intestinal metaplasia of the stomach is diagnosed histologically by the presence of acidic mucins within the epithelial cells and is associated with a higher risk of oncogenic progression. Incomplete metaplasia is further stratified into type II, which expresses sialylated mucins and type III, which express sulfated mucins within the epithelial cells (7, 8) with the sulfomucin expressing type III intestinal metaplasia being associated with the highest risk of progression to gastric adenocarcinoma (9-11). Similarly in the pancreas, expression of sulfated mucins occurs exclusively in high-risk lesions (Pancreatic intraepithelial neoplasia 3; PanIN-3) as well as pancreatic ductal adenocarcinoma(12-14). Sulfated mucins are also a feature of Barrett’s esophagus and esophageal adenocarcinoma (15). In sum, sulfated mucins are a high-risk feature of metaplasia throughout the gastrointestinal foregut.

Our group has shown that assaying for one of these sulfated epitopes (3’-Sulfo-Le^A/C^) using the antibody Das-1(16) is the most sensitive and specific means of diagnosing those pancreatic cysts harboring high-grade dysplasia and pancreatic ductal adenocarcinoma (12-14). The Das-1 antibody is also a highly sensitive and specific histological tool for Barrett’s esophagus (4, 15, 17, 18) and type III intestinal metaplasia of the stomach (4, 19, 20).

Coincident with the expression of these sulfated mucins in foregut metaplasia and cancer is the overexpression of specific galectins that preferentially bind these sulfated epitopes (21). Galectins are a family of lectins with affinity towards galactose containing N-Acetyllactosamine (LacNAc) moieties. Binding occurs between one or more of their carbohydrate recognition domains (CRD), which are a conserved fold formed by two, antiparallel, 5-6 strand beta-sheets that lie back-to-back. Galectin-3 is unique among the galectins in that it possesses a single CRD (22-28) and has a long proline rich N-terminal that tail with sequence similarity to collagen and which functions to multimerize galectin-3 and allow it to act as if contained multiple CRDs (29). Galectin-3 binds relatively weakly to either type I or type II terminal LacNAc moieties; however, the addition of a terminal sulfate on type II LacNAc significantly increased affinity (30, 31). Other acidic modifications like sialic acid did not similarly increase affinity(30).

The mouse stomach is uniquely suited for these studies because (1) unlike the human stomach, the murine stomach expresses sulfated mucins at homeostasis and (2) there exists widely used, synchronous, chemically induced experimental models to transform the stomach into spasmolytic polypeptide expressing metaplasia (SPEM). Here, we show that *in vivo* galectin-3 resides within the secretory zymogenic granules along with the sulfated mucins. This localization differs from the cytoplasmic distribution present *ex vivo* in cancer cell lines. Consistent with surveys of human tissue, both RNA and protein levels of galectin-3 increase as the murine zymogenic chief cells transition to metaplasia. We found that galectin-3 was necessary for selective cathartocytosis from organelles in which it resides, but not other cellular organelles. Inhibition of cellular downscaling manifests as a delay in the expression of metaplasia specific transcription factor SYR-box transcription factor 9 (Sox9) as well as proliferation.

## Results

### Galectin-3, Galectin-4, and the 20S proteosome bind Sulfomucins

To determine the repertoire of proteins that interact with secreted sulfomucins and might assist in their excretion during cathartocytosis, we performed immunoprecipitation of media conditioned by the LS174T cell line with the Das-1 antibody that is specifically reactive towards 3’-Sulfo-Le^A/C^. This cell line was chosen as it produces and secretes abundant sulfated mucins(32). Analysis with mass spectroscopy revealed a very small subset of proteins from the conditioned media (Figure 1A) that co-immunoprecipitated with sulfomucins. These included several components of the 20s proteosome, which will be investigated elsewhere, but relevant to this study also included two galectins (galectin-3 and galectin-4), which are known to preferentially associate with sulfated mucins relative to non-acidic glcyans and which are increased in gastric cancer relative to normal adjacent mucosa(21). We confirmed the mass spectroscopy results with western blot analysis (Figure 1B).

**Figure 1.**
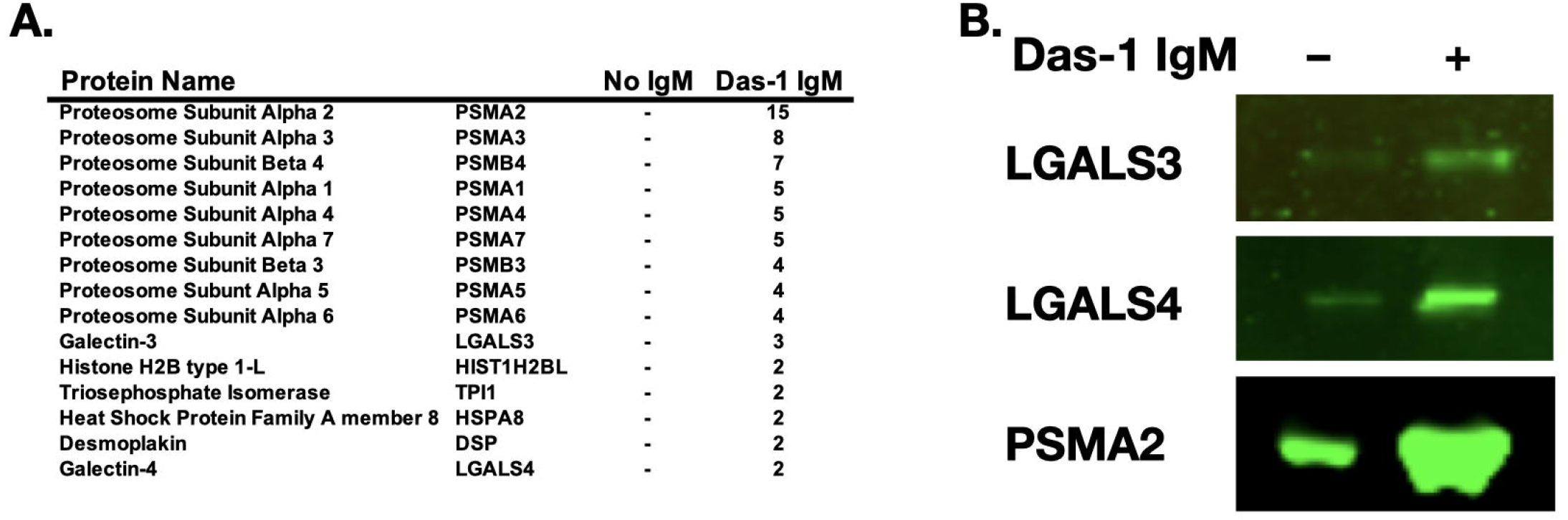

### Galectin-3, but not Galectin-4 Expression is Increased During the Transition to SPEM

We next measured RNA and protein levels of galectin-3 and galectin-4 both at baseline and during the transition to spasmolytic polypeptide expressing metaplasia (SPEM). Despite galectin-4 being increased in human gastric cancer relative to normal adjacent tissue, we did not observe a consistent increase in galectin-4 at either the RNA or protein level during the transition to SPEM (Supplemental Figure 1). These data suggest that galectin-4 might exert a putative effect much later in oncogenic cascade, which is supported by its ability to augment peritoneal metastases.(33) For these reasons, we will not consider galectin-4 further in this work.

In contrast to galectin-4, we found that expression of galectin-3 was significantly (~3-fold) increased at both the RNA and protein level (Figure 2A-C), suggesting at potential role in transition of normal, homeostatic tissue to SPEM.

**Figure 2.**
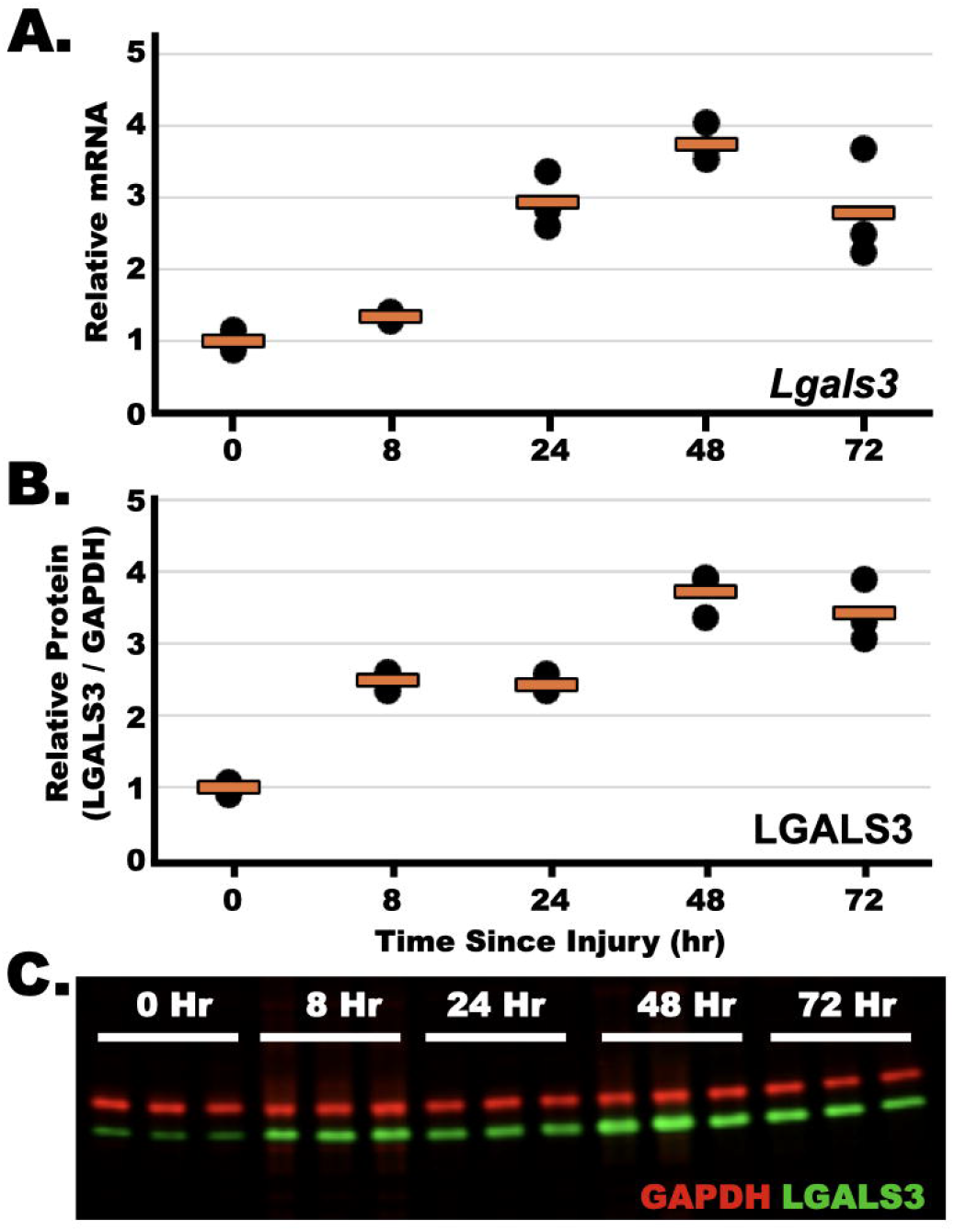

### Galectin-3 Co-Localizes with Sulfated Mucins in the Zymogenic Granules of the Chief Cell

Confocal microscopy demonstrated that galectin-3 colocalizes with sulfomucins within the zymogenic granules of the chief cell (Figure 3A). This *in vivo* localization differs from the *ex vivo* cytoplasmic and nuclear distribution in cancer cell lines, reported by others.(34) To ensure this effect was not due to a non-specific antibody, we show LGALS3 immunohistochemistry of galectin-3-sufficient C57BL/6J compared to galectin-3 knockout mouse. In C57BL/6J, we observe a vesicular distribution of galectin-3 in the chief cell and a homogeneous, cytoplasmic distribution in the pit cells (Figure 3B). No reactivity is observed in the stomach of the galectin-3 knockout mouse confirming the specificity of this antibody (Figure 3C).

**Figure 3.**
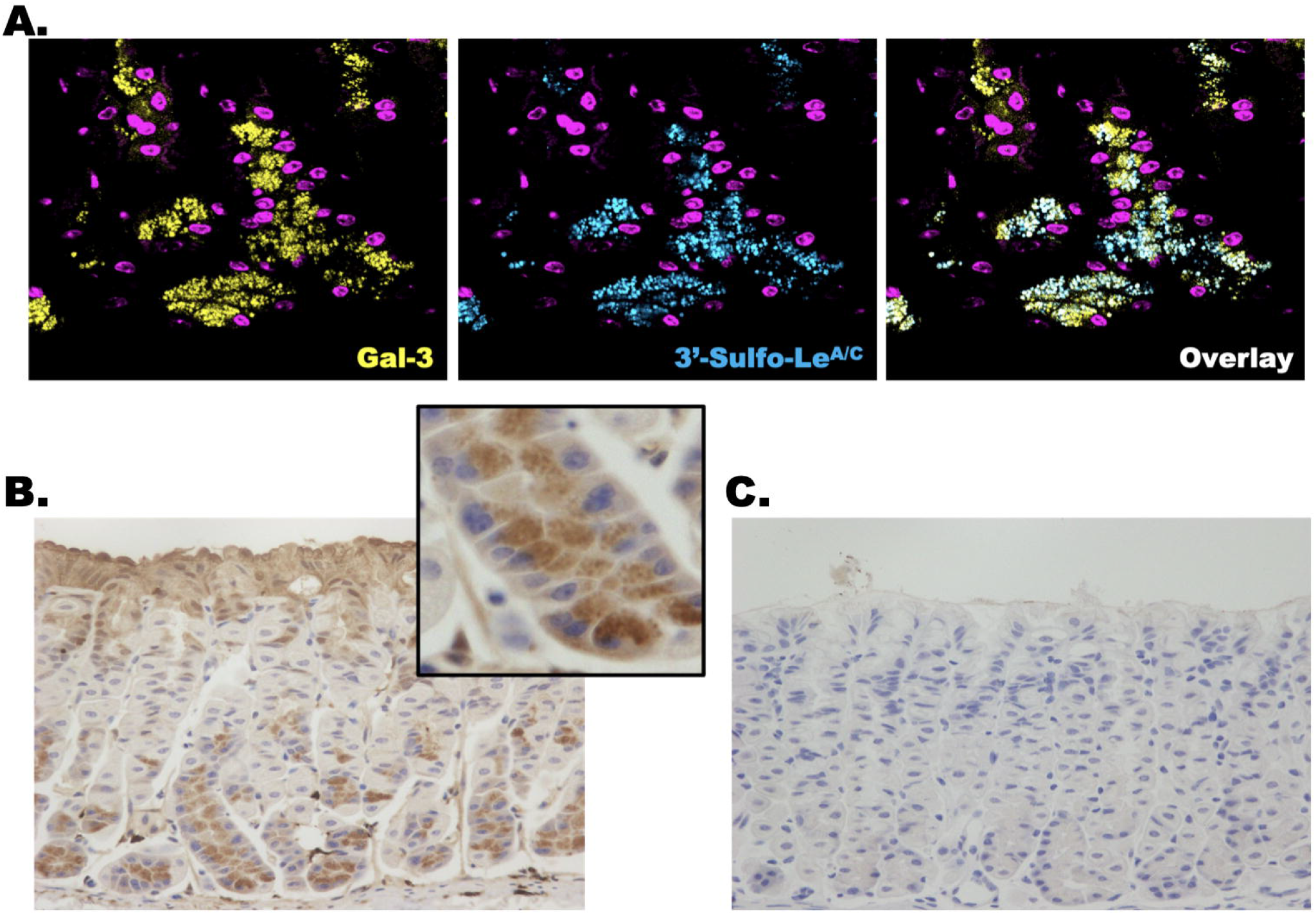

### Galectin-3 Aids in the Excretion of Sulfated Mucins during Cathartocytosis

Despite the presence of galectin-3 within the zymogenic chief cell granules at homeostasis, we could not tease out a significant histologic phenotype that differs between wild-type C57BL/6J mice and galectin-3 mice at homeostasis. However, during the metaplastic cascade induced chemically, we observed a dramatic defect in the dynamic secretion of sulfated mucins that occurs during the newly described process of cathartocytosis(35) (Figure 4B,C). Following injury, we observe retention of intracellular sulfomucins and a decreased number of gland lumens filled with sulfated mucins. In contrast to the zymogenic granules, where galectin-3 resides, we do not appreciate a difference in the excretion of endoplasmic reticulum as evidenced by equivalent percentage of gland lumens filled with Azure A reactive material. This suggests that galectin-3 is necessary for cathartocytotic excretion of the cellular compartments in which it resides but does not alter the excretion of other cellular compartments during cellular downscaling.

**Figure 4.**
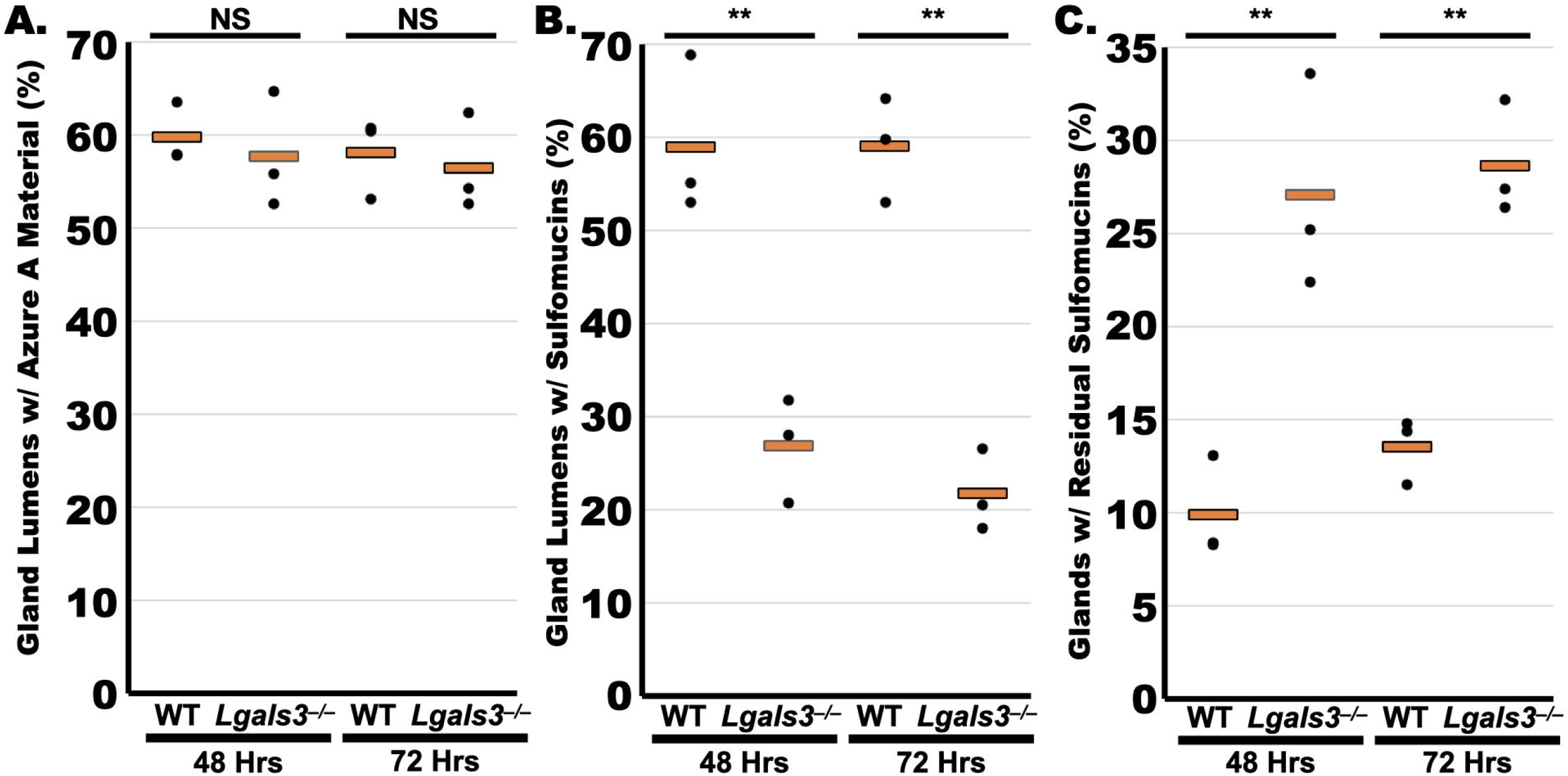

**Figure 5.**
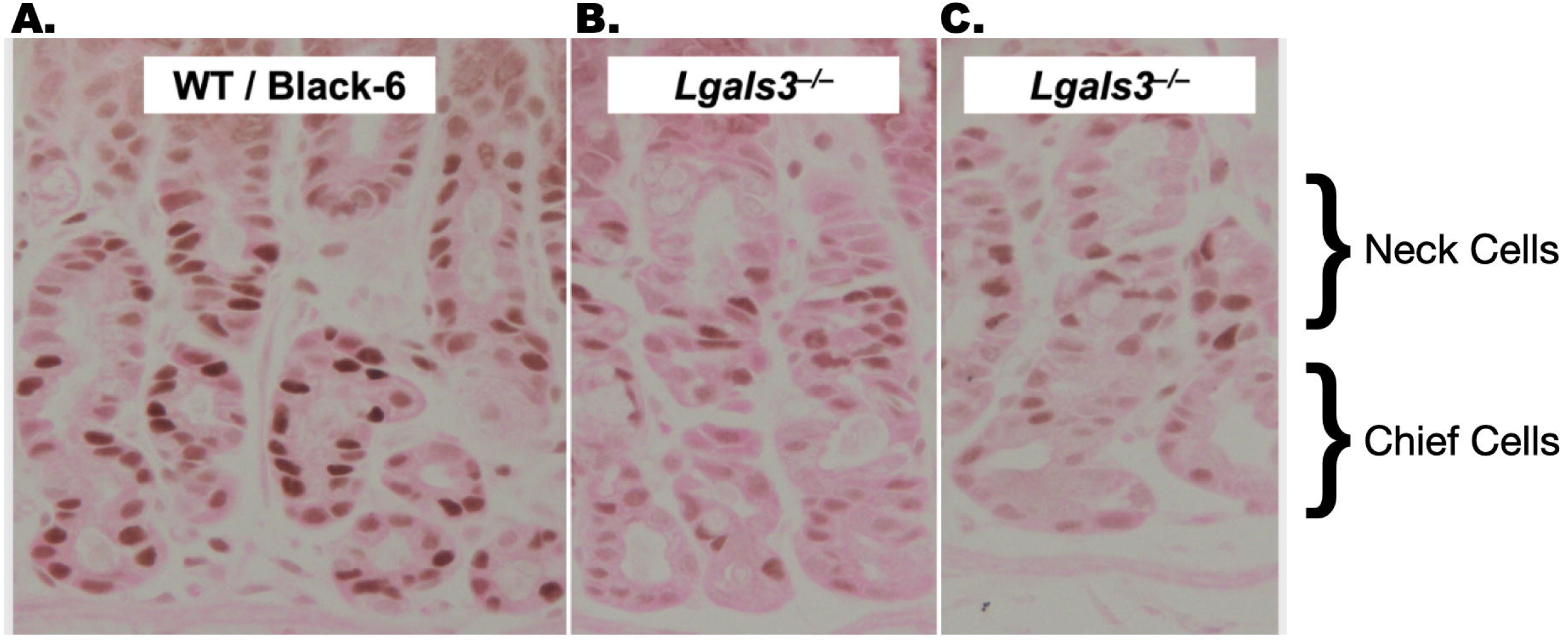

### Galectin-3 Knockout Impairs Expression of Transcription Factors and Proliferation

To investigate whether impaired cellular downscaling that we observe in *Lgals3*^*–/–*^ relative to wild-type C57BL/6J mice affected the metaplastic reaction, we examined the nuclear expression of Sox9, a transcription factor expressed in SPEM but not at homeostasis. In the galectin-3 null mice, we observe decreased expression of Sox9 in the base of the gland, the site where chief cells transition to SPEM (Figure 6). Neck cells, which also express Sox9 after injury, serve as an internal positive control. As Sox9 is expressed *en route* to proliferative metaplasia(36), we also measured proliferation. We found that the number of Ki-67 positive cells were significantly reduced in *Lgals3*^*–/–*^ relative to wild-type C56BL/6J and that the defect was most prominent at the bases of the gland.

**Figure 6.**
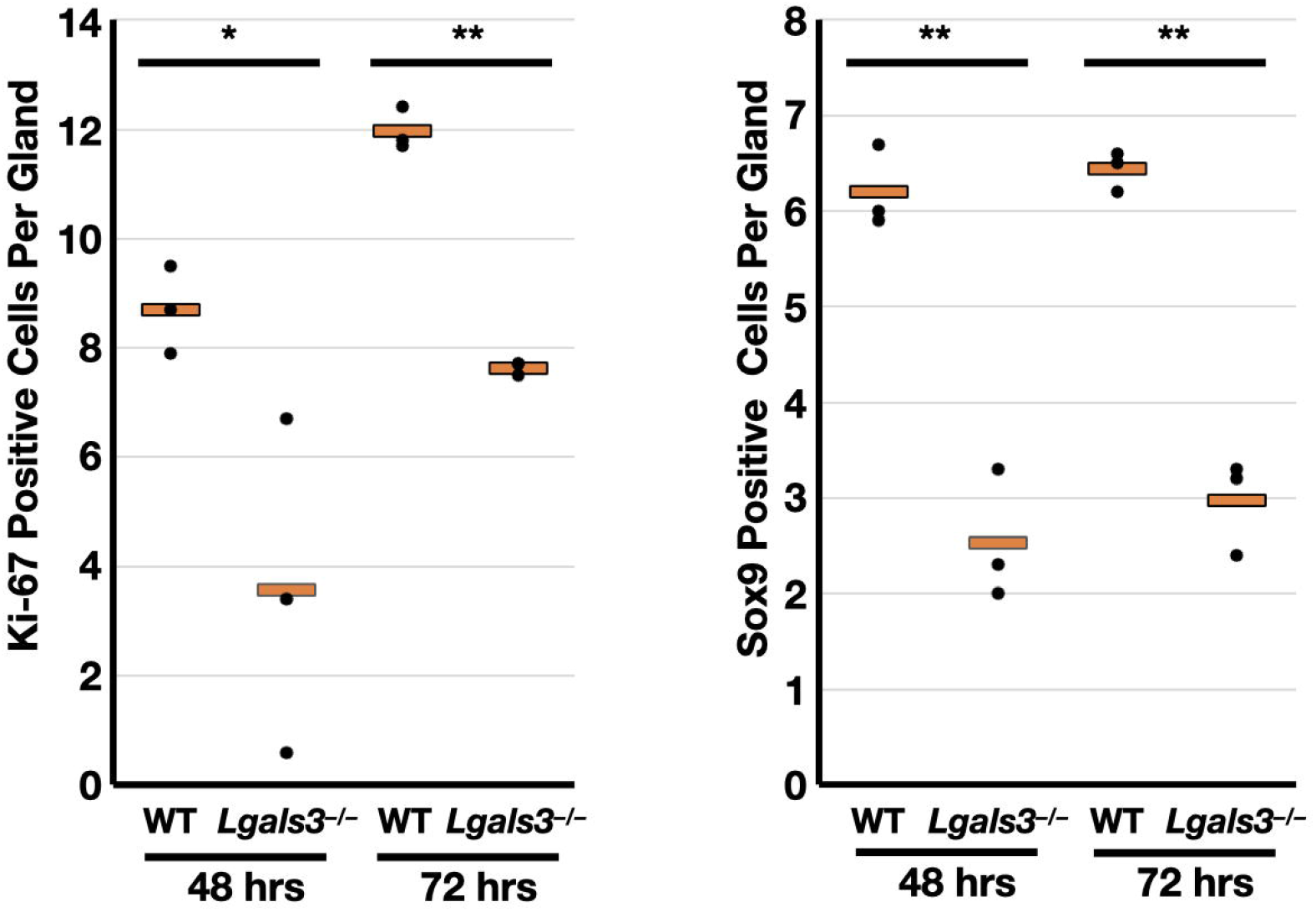

## DISCUSSION

Following injury, tissue must adapt to replace lost and irreparably damaged cells to restore tissue homeostasis. For cell-types derived from a stem cell and for which the stem cell population is still viable this can be accomplished though proliferation and subsequent differentiation (e.g. small intestine). However, in many tissues and for several cell-types no such stem cell exists and therefore regeneration relies on plasticity of “terminally-differentiated” cells. Both pulse-chase DNA labeling experiments (37) as well as genetic lineage tracing(38) have demonstrated that zymogenic chief cells are such a cell population that are responsible for their own census and do not derive from the isthmal stem cell / transit amplifying population.

The stepwise process where mature chief cells transform into a dedifferentiated proliferative cell to replace lost cells, prior to redifferentiation into a mature chief cell is called paligenosis.(39-41) In the original description of paligenosis, it was assumed that the cellular downscaling necessary to produce a proliferative cell from a mature cell was accomplished exclusively by canonical autophagy.(39, 42) However, we recently described a parallel process, cathartocytosis, that occurs concurrently with canonical autophagy during paligenosis, but is mechanistically independent.(35)

Here, we identify galectin-3 as first [of presumably many] protein that participates in cathartocytosis. Unlike the *ex vivo* intracellular distribution in cancer cell lines, we found that galectin-3 was specifically localized in the zymogenic granules of chief cells at homeostasis. Following injury, we found that cathartocytotic excretion of this cellular compartment was compromised; however, other cellular organelles like the endoplasmic reticulum that lack galectin-3 were not impacted. Such a result, is consistent with galectin-3 organizing either directly or indirectly protein tethering and fusing complexes on the vesicle membranes in which it resides. For example, another galectin, galectin-9 has been shown to bind to VAMP-3, a vesicular soluble NSF attachment protein receptor (v-SNARE) and alter vesicular trafficking.(43) Galectin-3 shown to interact with components of the SNARE complexes in other contexts.(44) Future studies, will elucidate precisely how galectin-3 participates in cathartocytotic excretion.

Our data also demonstrates that cathartocytosis is necessary for the paligenotic progression to proliferative SPEM. The blockade of the cathartocytotic excretion of the zymogenic granules in *Lgals3*^*–/–*^ mice led to repression of SOX9 expression and inhibition of proliferation in SPEM. Such a finding validates (1) the importance of cathartocytosis in the metaplastic reaction and (2) the necessity to downscale cellular machinery prior to licensing a cell to proliferate in paligenosis.

Lastly, the mechanism by which galectins (and other proteins like annexins) are secreted has been a matter of debate.(45) It has been established that these two classes of proteins are generally not secreted via canonical pathways; however, the mechanism by which they are secreted is unknown. Our data, suggests that cathartocytosis may represent one such way these proteins are secreted. We discovered cathartocytosis by studying how cells downscale their cellular machinery in response to injury. This post-injury cellular reaction is consistent with numerous reports that serum galectin-3 levels are a biomarker prognostic marker for innumerable cellular injuries like myocardial infarction(46), heart failure(47), hemorrhagic stroke(48), kidney disease(49), autoimmune processes(50-52), cancers(53). Such widespread expression and secretion of galectin-3 suggests that cathartocytosis and/or other unconventional secretory mechanisms are common cellular responses to tissue injury in widespread clinical contexts.

## Materials & Methods

### Animal studies and reagents

All experiments using animals followed protocols approved by the Washington University in St. Louis, School of Medicine Institutional Animal Care and Use Committee. Both wild-type C57BL/6 as well as a breeding pair of Galectin-3 knockout mice (*B6*.*Cg-Lgals3*^*tm1Poi*^*/J*)(54) were purchased from Jackson laboratories (Bar Harbor, ME). Galectin-3 knockout mice were bred homozygous KO x homozygous KO.

Tamoxifen powder (Toronto Research Chemicals) was initially solubilized in 100% ethanol via sonification after which it was emulsified in sunflower oil (Sigma-Aldrich) at a 9 Oil :1 EtOH ratio(55). Tamoxifen (5 mg/20 g body weight; Toronto Research Chemicals) was injected intraperitoneally daily for up to 2 days or until mouse was euthanized for histologic examination. Hydroxychloroquine (120 mg/Kg by intraperitoneal injection) was administered 24 hours prior to the first tamoxifen injection and then with all subsequent tamoxifen injections until the tissue was harvested. All mouse experiments were performed on mice aged 6-10 weeks.

### Imaging and tissue analysis

Following anesthetizing with isoflurane and cervical dislocation, murine stomachs were excised, flushed with phosphate buffered saline and fixed overnight with 10% formalin. They were washed and equilibrated in 70% ethanol for several hours prior to embedding in 3% agar and routine paraffin processing. Sections (5-7 μM) were prepared for immunohistochemistry and/or immunofluorescence by deparaffinization using Histoclear and an alcohol series for rehydration. After which antigen retrieval was performed in 10 mM citrate buffer, pH 6.0 in a pressure cooker. Tissue was blocked with 2% BSA and 0.05% Triton X-100. Primary and secondary antibodies were diluted in 2% BSA and 0.05% Triton X-100. Vector ABC Elite kit was used for immunohistochemistry. Immunohistochemistry slides were mounted with Permount and immunofluorescence with Prolong Gold. Brightfield images were taken on either a Nanozoomer (Hamamatsu 2.0-HT System) for quantitation or Olympus BX43 light microscope. Confocal images were obtained on a Zeiss LSM880 confocal microscope.

### Azure A staining

Slides of paraffin embedded tissue were dewaxed and rehydrated with an ethanol series to 30%. Antigen retrieval must not be performed as this results in loss of staining. Slides were then stained with 0.1% Azure A Chloride in 30% ethanol for 1 minute. Slides were then washed in water. Placing slides in citric acid / sodium phosphate buffer, pH 4 results in royal blue nuclear staining and dark blue cytoplasmic staining. The slides were then transferred to 95% ethanol and stained with Eosin Y. The slides were subsequently mounted with permount.

### Western blot

Western blot samples were prepared using ~30 μg of protein in standard SDS-PAGE lammeli buffer with 5% β-mercaptoethanol. Samples were heated to 70 ºC for 10 minutes prior to running through NuPAGE precast gels. Protein was transferred in NuPAGE transfer buffer containing 20% methanol to nitrocellulose which was blocked using 5% BSA in PBS. Nitrocellulose membrane was incubated with primary antibodies in blocking buffer overnight at 4 ºC. Membranes were washed with PBS and then incubated in appropriate secondary antibodies at 1:10,000 dilution in blocking buffer. After extensively washing, membrane was imaged using LiCOR Odyssey.

### Fundic Chief Cell Size

2-Dimensional area measurements of chief cells were made by analyzing 40x micrographs of hematoxylin and eosin stained sections. Cells were outlined manually and area calculated with ImageJ/FIJI(56). 270 pixels / 50 μm. Only well-defined chief cells within 3 cells of the base of gland with both nuclei and luminal apex present were included in the calculations.

#### List of Materials

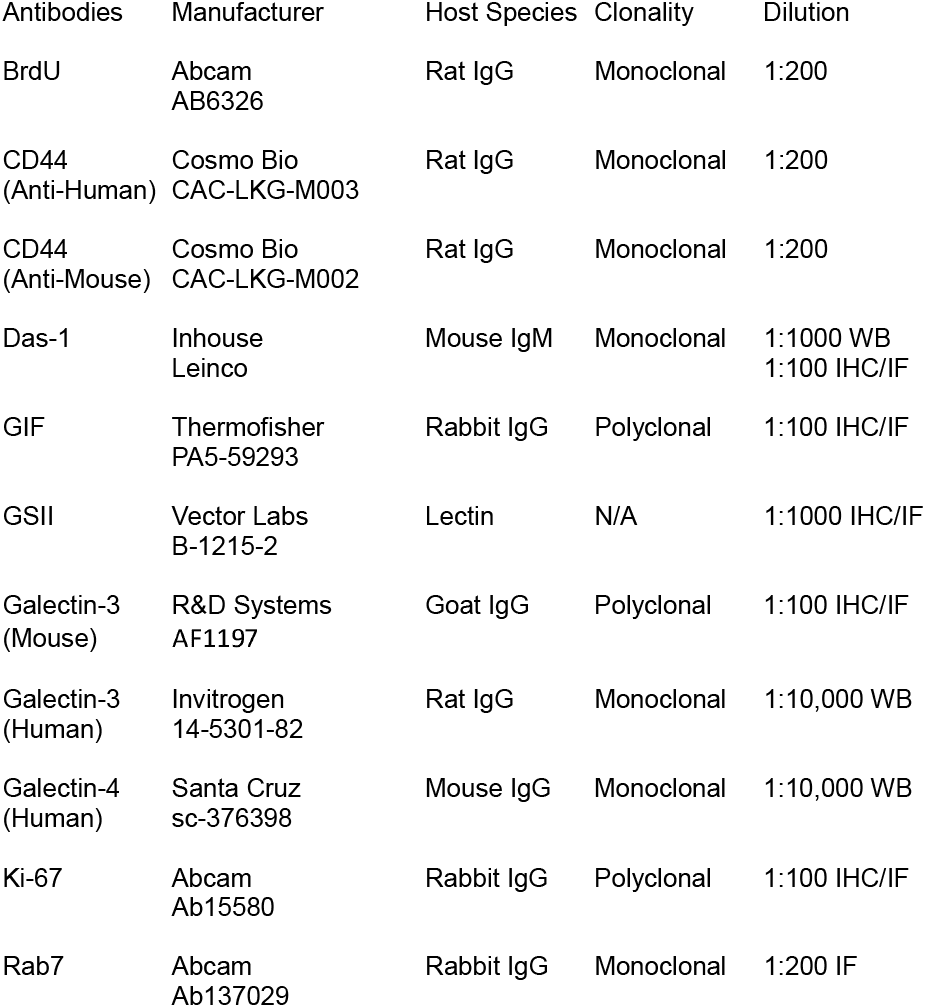
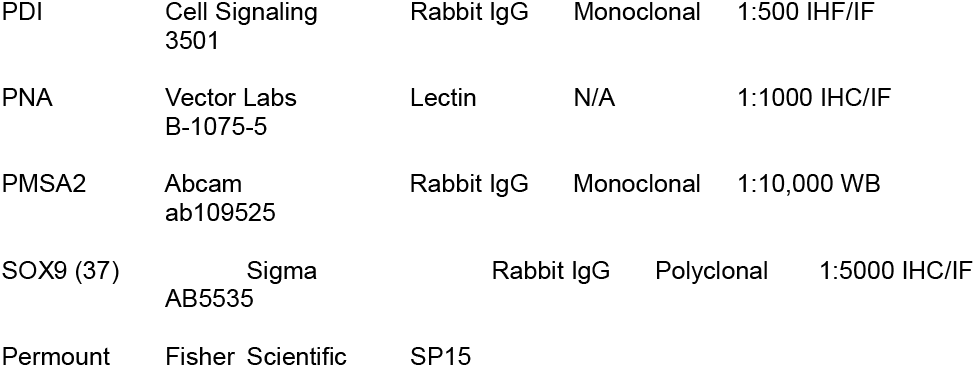

## ACKNOWLEDGEMENTS

Jeffrey W. Brown is K08 DK132496; P30 DK052574; The Foundation for Barnes-Jewish Hospital

## Author Contributions

Study Concept and Design: XL, JWB. Data Acquisition and Analysis: XL, GN, XL, AP, MH, JWB; Initial draft of the manuscript: JWB; Revisions to the manuscript: XL, GN, XL, JWB

---

**Supplemental Figure 1.**
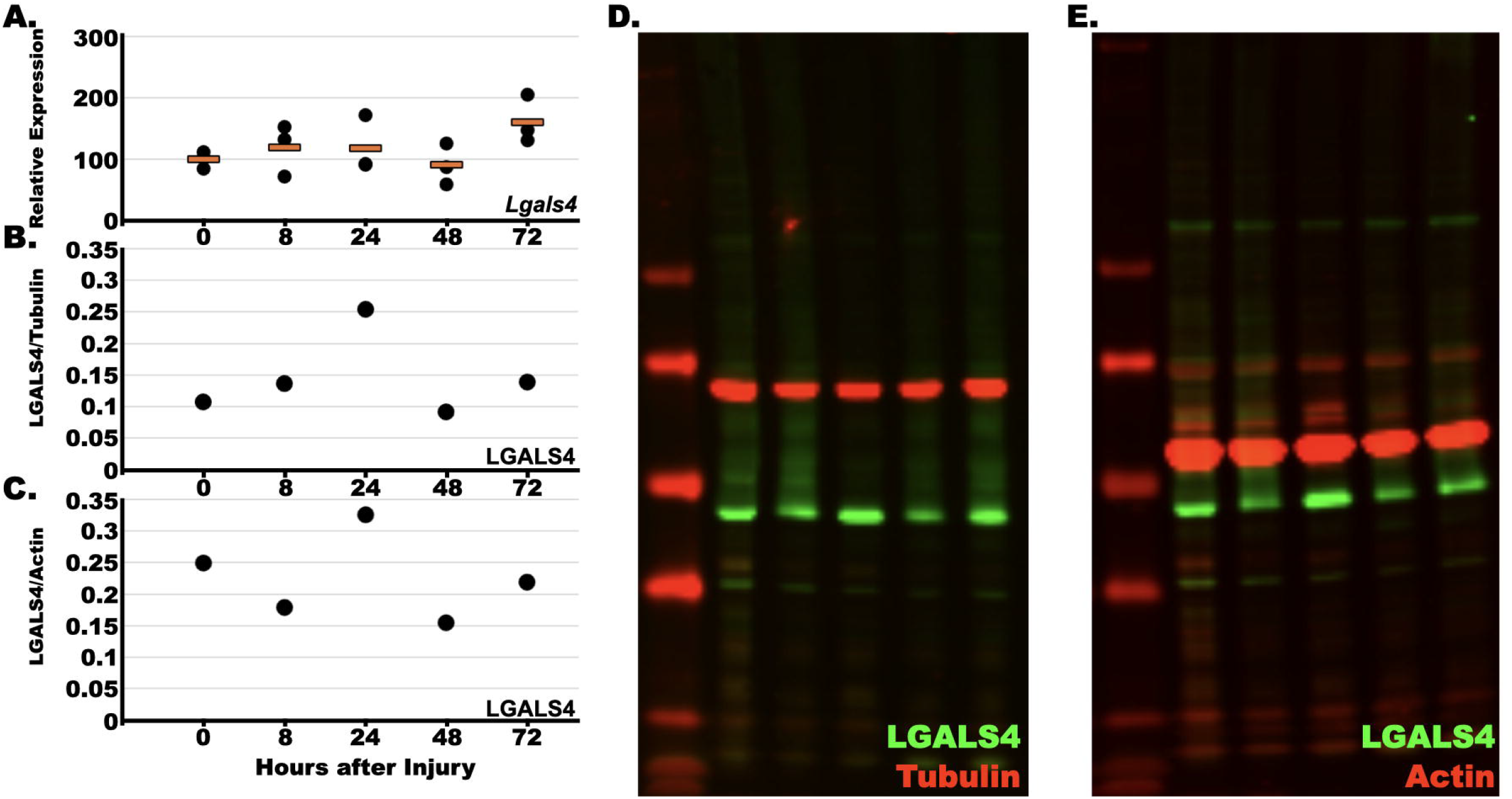

